# Low- and high-grade glioma endothelial cells differentially regulate tumor growth

**DOI:** 10.1101/2023.07.07.548125

**Authors:** Sree Deepthi Muthukrishnan, Haocheng Qi, David Wang, Lubayna Elahi, Amy Pham, Alvaro G. Alvarado, Tie Li, Fuying Gao, Riki Kawaguchi, Albert Lai, Harley I. Kornblum

## Abstract

**Background:** A key feature distinguishing high-grade glioma (HGG) from low-grade glioma (LGG) is the extensive neovascularization and endothelial hyperproliferation. Prior work has shown that tumor endothelial cells (TEC) from HGG are molecularly and functionally distinct from normal brain EC and secrete higher levels of pro-tumorigenic factors that promote glioma growth and progression. However, it remains unclear whether TEC from LGG also express pro-tumorigenic factors, and to what extent they functionally contribute to glioma growth.

**Methods:** Transcriptomic profiling was conducted on tumor endothelial cells (TEC) from grade II/III (LGG, IDH-mutant) and grade IV HGG (IDH-wildtype). Functional differences between LGG- and HGG-TEC were evaluated using growth assays, resistance to anti-angiogenic drugs and radiation therapy. Conditioned media and specific factors from LGG- and HGG-TEC were tested on patient-derived gliomasphere lines using growth assays *in vitro* and in co-transplantation studies *in vivo* in orthotopic xenograft models.

**Results:** LGG-TEC showed enrichment of extracellular matrix and cell cycle-related gene sets and sensitivity to anti-angiogenic therapy whereas HGG-TEC displayed an increase in immune response-related gene sets and anti-angiogenic resistance. LGG- and HGG-TEC displayed opposing effects on growth and proliferation of IDH-wildtype and mutant tumor cells. Asporin (ASPN), a small leucine rich proteoglycan enriched in LGG-TEC was identified as a growth suppressor of IDH-wildtype GBM by modulating TGFΒ1-GPM6A signaling.

**Conclusions:** Our findings indicate that TEC from LGG and HGG are molecularly and functionally heterogeneous and differentially regulate the growth of IDH-wildtype and mutant tumors.

## Introduction

High-grade gliomas (HGG, Grade IV) are more extensively vascularized than low-grade gliomas (LGG, Grade II/III), with endothelial hyperproliferation serving as a key histopathological hallmark differentiating these tumors^1^. Despite being highly angiogenic tumors, anti-angiogenic therapies have largely been unsuccessful in impeding tumor growth or improving patient survival outcomes in HGG. This resistance is mainly due to activation of alternative neovascularization mechanisms such as vessel co-option, vascular mimicry, vasculogenesis and endothelial transdifferentiation and activation of other pro-angiogenic pathways ^2, 3^. Nevertheless, the neoplastic vessels generated by these mechanisms are highly dysfunctional, leaky, and disorganized. Prior studies including our work have demonstrated that tumor endothelial cells (TEC) from HGG are molecularly heterogeneous compared to normal brain endothelial cells (NEC)^4–9^.

The vasculature of LGG is not well studied compared to HGG, and it remains unclear whether there is molecular heterogeneity in TEC from different LGG. Importantly, the majority of adult LGG have mutations in the enzyme Isocitrate dehydrogenase (IDH)1 or IDH2 with 50-80% reported in Grade II and 54% in Grade III gliomas^10^. On the contrary, only 15-20% of Grade IV gliomas harbor mutations in IDH1 or IDH2, indicating that IDH mutation status may govern the vascular phenotype, and this could in turn influence their sensitivity to anti-angiogenic therapies. A recent study reported key differences in angiogenic gene expression related to hypoxia and TGFβ signaling between LGG (Grade II) IDH-wildtype and mutant tumor vessels^11^. It remains undetermined to what extent the molecular landscape of TEC from LGG differ from HGG, and how it influences their response to anti-angiogenic treatments, and whether the angiocrines expressed in LGG-TEC exhibit pro-tumorigenic functions.

In this study, we conducted transcriptomic profiling of TEC isolated and cultured from IDH-mutant (mIDH) Grade II/III LGG and IDH-wildtype (wIDH) Grade IV HGG that included primary and recurrent tumors. We show that TEC from LGG and HGG exhibit significant molecular and functional heterogeneity and differential sensitivity to anti-angiogenic therapy. LGG- and HGG- TEC differentially regulate the growth of wIDH GBM *in vitro* and in orthotopic xenograft models. Differential gene expression analysis revealed several extracellular matrix proteins are enriched in LGG-TEC and inflammatory cytokines and chemokines in HGG-TEC. Specifically, we identified ASPORIN (ASPN), a member of the small leucine rich proteoglycan family, highly enriched in LGG-TEC as a potential tumor suppressor that differentially regulates the growth of wIDH and mIDH tumors via modulation of TGFβ1 signaling.

## Materials and Methods

1. **Patient-derived gliomasphere lines:** All patient-derived gliomasphere lines utilized in this study were previously established in our laboratory. Gliomaspheres were cultured in DMEM/F12 medium supplemented with B27, 20 ng/ml bFGF, 50ng/ml EGF, 5μg/ml Heparin, and antibiotics penicillin/streptomycin. Gliomaspheres were dissociated into single cells every 7-14 days depending on growth rate, and experiments were performed with cell lines that were cultured for < 20 passages since their initial establishment, and tested negative for mycoplasma contamination. Cell lines were authenticated by STR analysis.
2. **Culture of tumor endothelial cells and human brain endothelial cells:** TEC (P1 to P9) and HBEC (Sciencell) were cultured in endothelial cell growth media (ECM) (R&D systems) in tissue culture flasks. Validation of endothelial identity was done using CD31 immunostaining at P2 and P7 after expansion. Detailed protocol for isolation of TEC from patient tissue is provided in supplementary methods.
3. **RNA sequencing and analysis:** Bulk-RNA sequencing and single-cell RNA sequencing dataset analysis was carried out as described previously^8, 12^.
4. **Animal strains, intracranial transplantation and imaging**: All animal studies were performed according to approved protocols by the institutional animal care and use committee at UCLA. Studies did not discriminate sex, and both male and females were used. **Strains**: 10-12-week old NOD-SCID gamma null (NSG) mice were used to generate orthotopic xenografts. 5X10^4^ cells from a patient-derived GBM line (HK_408) containing a firefly-luciferase-GFP lentiviral construct were injected intracranially into the neostriatum in mice. Co-transplantation with TEC- expressing mCherry was performed at a ratio of 1:1 (tumor: endothelial cells), with 5X10^4^ cells per condition. **Imaging**: Tumor growth was monitored 2 and 4 weeks after transplantation by measuring luciferase activity using IVIS Lumina II bioluminescence imaging. ROIs were selected to encompass the tumor area and radiance was used as a measure of tumor burden.
5. **Lentiviral constructs and gene knockdown**: PLV-mCherry (Vector builder), shRNA-scrambled, shRNA-ASPN-GFP and shRNA-GPM6A-GFP (abmgood) were purchased from manufacturers as indicated. Briefly, cells were transduced with the lentivirus and reporter expression was analyzed at 48 hours. Following reporter activity, cells were selected with Puromycin (Sigma Aldrich) for 72 hours and knockdown of respective genes was confirmed by quantitative RT-PCR and western blotting.
6. **Statistical analysis.** All data are expressed as the mean ± SD. *P* values were calculated in Graph Pad Prism 8.0 using unpaired two-tailed Student t-test and ANOVA for multiple comparison followed by bonferroni correction and post hoc t-test. *P* values of less than 0.05 were considered significant. Log-rank analysis was used to determine the significance of Kaplan-Meier survival curves. All other materials and methods are described in the supplemental information.

## Results

### Tumor endothelial cells are molecularly distinct from normal brain endothelial cells

To determine whether tumor endothelial cells (TEC) exhibit molecular heterogeneity, we established CD31+ TEC cultures isolated from glioma patients. TEC identity was validated by immunostaining and quantitative RT-PCR of endothelial markers (Figure 1A, B). RNA-sequencing and differential gene expression analysis (DEA) of glioma and TEC fractions from a primary glioblastoma (GBM) showed that the tumor fraction was enriched with glioma-stem cell (GSC) markers, and TEC fraction significantly expressed endothelial genes but not GSC markers, confirming their respective identities (Figure 1C).

**Figure 1:**
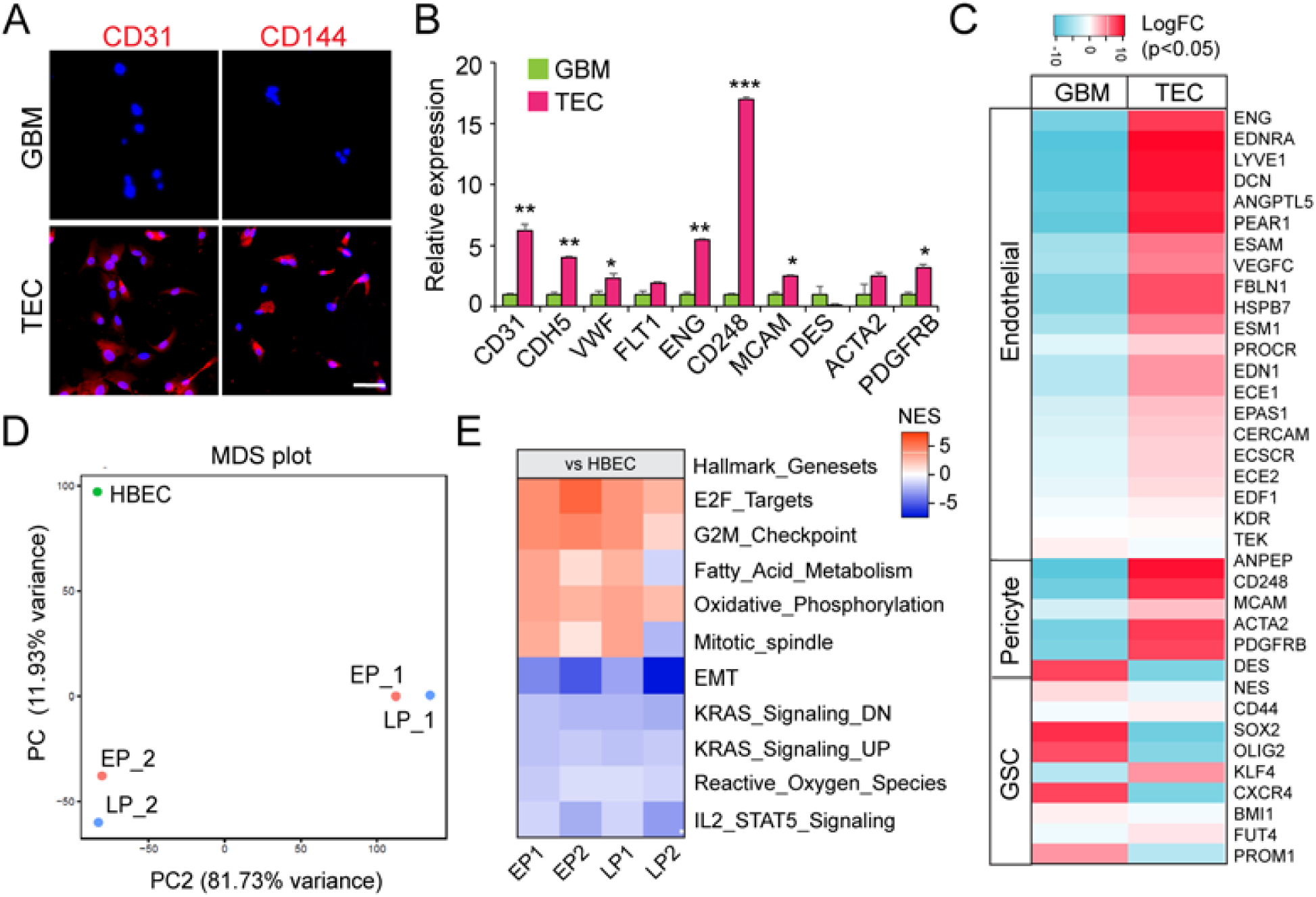
Molecular differences between normal brain-and tumor-endothelial cells. **A.** Immunostaining of CD31 (red) and CD144 (VE-CADHERIN, red) and DAPI (nuclei, blue) in patient-derived GBM and TEC fractions. Scale bars, 100μm. **B.** Relative expression of endothelial and pericyte markers in patient-derived GBM and TEC fractions. N=3, * p<0.05, ** p<0.005 and *** p<0.0005, unpaired t-test. **C.** Heatmap of LogFc expression of endothelial, pericyte and glioma stem cell (GSC) markers in GBM and TEC fractions. **D.** MDS plot of TEC cultured from early (EP) and late (LP) passages and HBEC. **E.** Heatmap of gene sets enriched in early and late passage TEC compared to HBEC.

Next, to ensure that TEC maintained their endothelial identity in long-term culture, we performed immunostaining for CD31 at early (P1-P2) and late (P5-P7) passages, and found that the expression was maintained over several passages (Figure S1A). Further, we performed RNA-sequencing on early (P1-P2) and late (P4-P7) passage TEC isolated from two GBM patients and compared to cultured human brain microvascular endothelial cells (HBEC). Principal component analysis (PCA) showed that TEC and HBEC clustered separately indicating that they are molecularly distinct (Figure 1D). DEA also showed that cultured TEC are significantly distinct from HBEC, but there is minimal difference between early and late passage TEC cultures (Figure S1B). Gene set enrichment analysis (GSEA) revealed enrichment of cell cycle and DNA-repair related processes in TEC compared to HBEC (Figure 1E). We also compared the gene expression profiles of cultured and freshly isolated TEC and found that they closely clustered together (Figure S1C). These data strongly indicate that TEC are molecularly distinct from HBEC.

### LGG- and HGG-TEC display molecular and functional heterogeneity

Given that microvascular proliferation is a distinguishing feature of Grade IV GBM, we wondered if TEC from LGG and HGG exhibited molecular and functional heterogeneity^2^. To test this, we performed whole transcriptomic sequencing of TEC cultures from Grade II/III, mIDH LGG (N=5) and Grade IV, wIDH primary (N=4) and recurrent GBM (N=5). Differential gene expression analysis showed that TEC from recurrent (REC) GBM are highly distinct from LGG-TEC but exhibit minimal differences in comparison with primary (PRI)-TEC (Figure 2A, B). GSEA showed that HGG-TEC are significantly enriched for immune response related gene sets, whereas LGG- TEC are enriched for cell-cycle related processes.

**Figure 2:**
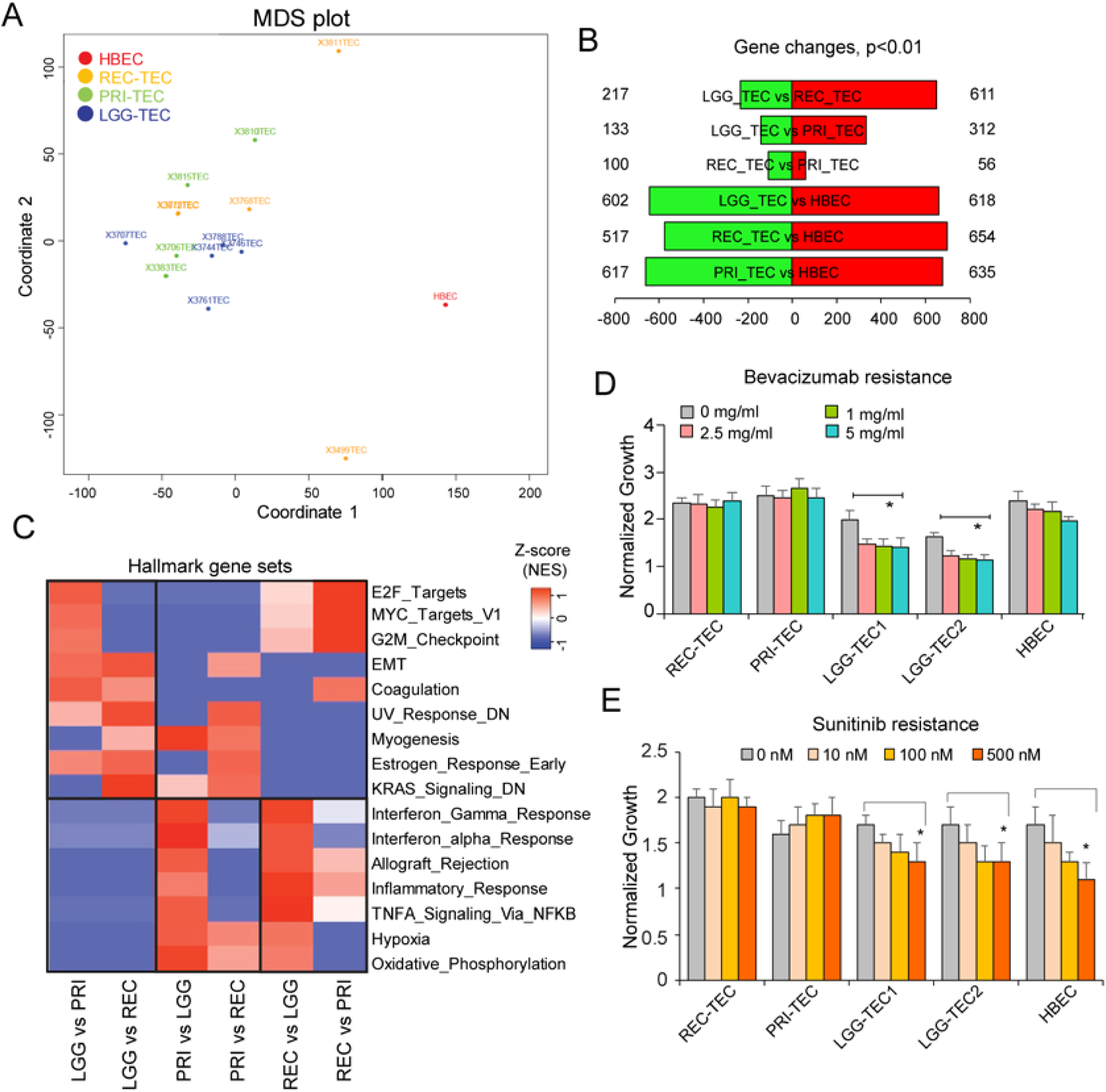
Molecular and functional heterogeneity of LGG- and HGG-TEC. **A.** MDS plot of LGG-, HGG (primary/PRI and recurrent/REC)-TEC and HBEC **B.** Genes differentially expressed between LGG-TEC, HGG-TEC and HBEC. **C.** Heatmap of normalized enrichment scores of gene sets enriched in LGG-TEC, PRI-TEC and REC-TEC. **D.** Normalized growth of LGG-TEC, PRI-TEC, REC-TEC and HBEC cultured in different doses of Bevacizumab and Sunitinib. N=3, * p<0.05, one-way ANOVA, post-hoc t-test.

Consistent with prior findings, we found that both LGG-TEC and HGG-TEC are morphologically distinct from HBEC, and displayed different rates of proliferation as assessed by EdU incorporation in culture^4, 5, 11^ (Figure S2A, B). Both LGG- and HGG-TEC also showed greater migration capacity than HBEC (Figure S2C). HBEC and LGG-TEC displayed higher sensitivity to anti-angiogenic treatments including Bevacizumab and Sunitinib, whereas HGG-TEC were resistant even at higher doses (Figure 2D, E). Furthermore, TEC were highly resistant to high-doses of radiation (8-10 Gy) compared to HBEC (Figure S2D). These findings indicate HGG-TEC exhibit greater capacity for treatment-resistance than LGG-TEC, and they are molecularly and functionally distinct from HBEC.

### LGG-TEC and HGG-TEC differentially regulate the growth of wIDH GBM and mIDH astrocytoma

Based on the molecular and functional differences between LGG- and HGG-TEC, we postulated that they may differentially influence the growth of tumor cells. We therefore collected conditioned media (CM) from LGG-, PRI- and REC-TEC cultures to determine if they differentially regulate the growth of GBM tumor cells. First, we tested the effects of TEC-CM on wIDH GBM lines (408, 301 and 336) and found that HGG-TEC promoted, whereas LGG-TEC significantly inhibited the growth of these tumor lines (Figure 3A, B and Figure S3A). Surprisingly, HEK293T cells used as a control showed significant growth inhibition on all the GBM lines. On the contrary, HGG-TEC did not alter the growth of mIDH astrocytoma line, whereas LGG-TEC slightly promoted their growth, indicating that they differentially affect the growth of wIDH GBM and mIDH astrocytoma tumor cells (Figure 3C, Figure S3B).

**Figure 3:**
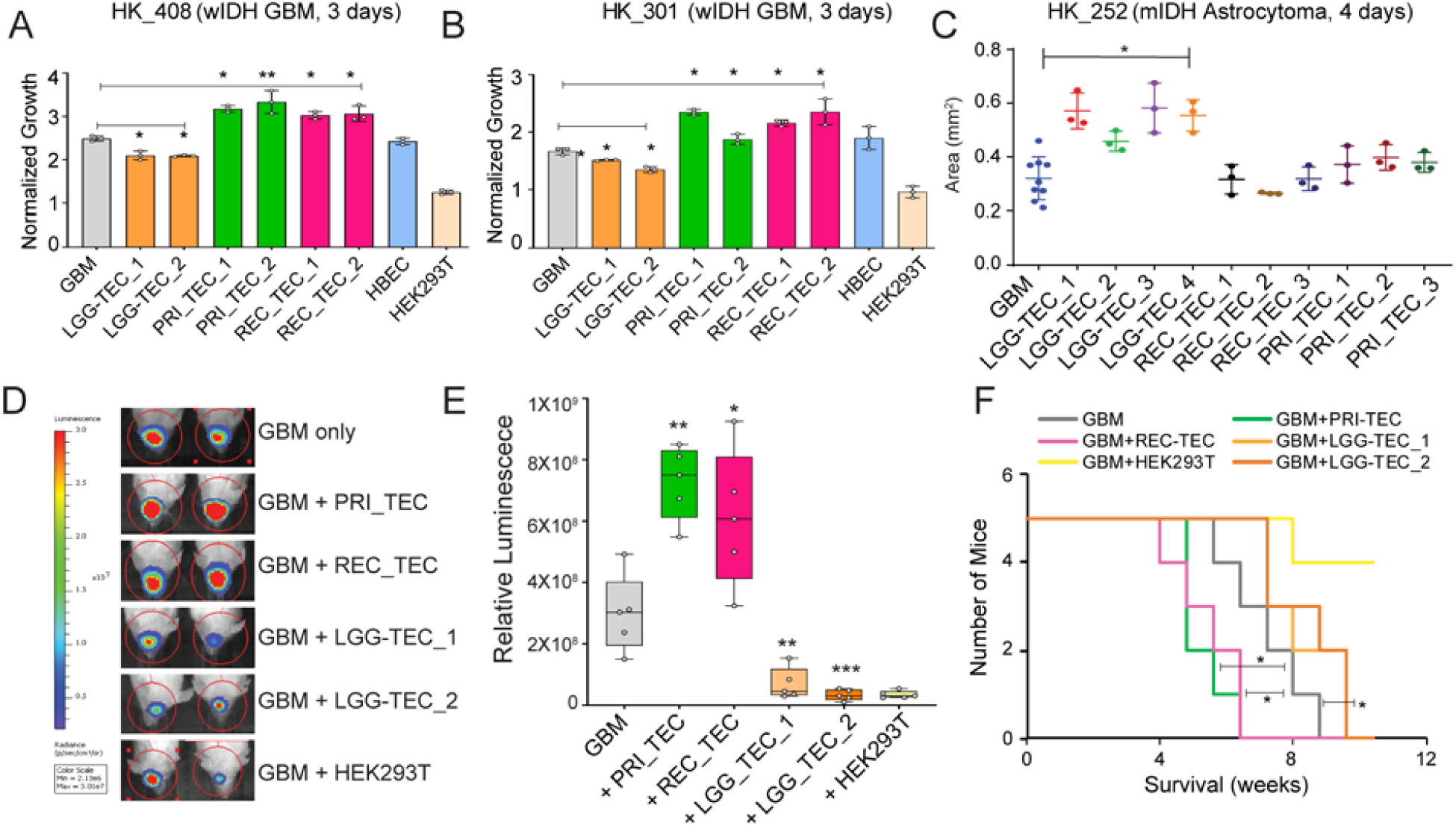
LGG-TEC and HGG-TEC differentially regulate the growth of GBM tumors. **A.** Normalized growth of wIDH GBM lines cultured in conditioned media from LGG_TEC, PRI_TEC and REC_TEC. HBEC was used for normal brain endothelial cells, and HEK293T cells were used as a negative control. * p<0.05, ** p<0.005, one-way ANOVA. **B.** Area of mIDH astrocytoma spheroids in conditioned media from LGG_TEC, PRI_TEC and REC_TEC. N=3 replicates per condition. * p<0.05, one-way ANOVA. **C.** Representative bioluminescent images of tumor growth of wIDH GBM cells co-transplanted with PRI_TEC, REC_TEC, LGG_TEC or HEK293T. Box plots of relative luminescence from tumors in each condition. N=5 mice per group. * p<0.05, ** p<0.005 and *** p<0.0005, one-way ANOVA and post-hoc t-test. **D.** Kaplan-Meier survival curve of mice co-transplanted with GBM and TEC. * p<0.05, Log-rank test.

To validate the *in vitro* findings, we co-transplanted either LGG-TEC or HGG-TEC along with wIDH GBM cells expressing firefly-luciferase-GFP into immunocompromised mice to generate orthotopic xenografts. Examination of tumors 4-weeks post-transplantation showed that LGG-TEC significantly inhibited the growth of the tumor cells, whereas both PRI- and REC-TEC enhanced the growth of the tumors (Figure 3D, E). In line with the *in vitro* data, HEK293T cells inhibited the growth of GBM tumors. This was also reflected in animal survival, as tumors co-transplanted with HEK293T or LGG-TEC significantly survived longer, and mice bearing tumors with HGG-TEC showed significantly reduced survival (Figure 3F). These findings strongly suggest that LGG- and HGG-TEC differentially regulate wIDH GBM growth.

### LGG-TEC and HGG-TEC show differential expression of extracellular matrix proteins, growth factors and cytokines

To elucidate the mechanism underlying the differential effects of TEC on tumor growth, we examined the transcriptomic data for secreted factors differentially expressed between LGG-vs HGG-TEC. Interestingly, we found several extracellular matrix proteins enriched in LGG-TEC that were either not expressed or showed minimal expression in HGG-TEC. Similarly, we found increased expression of chemokines and cytokines in HGG-TEC that showed virtually little to no expression in LGG-TEC. We also verified that these transcripts were enriched in TEC relative to HBEC (Figure 4A).

**Figure 4:**
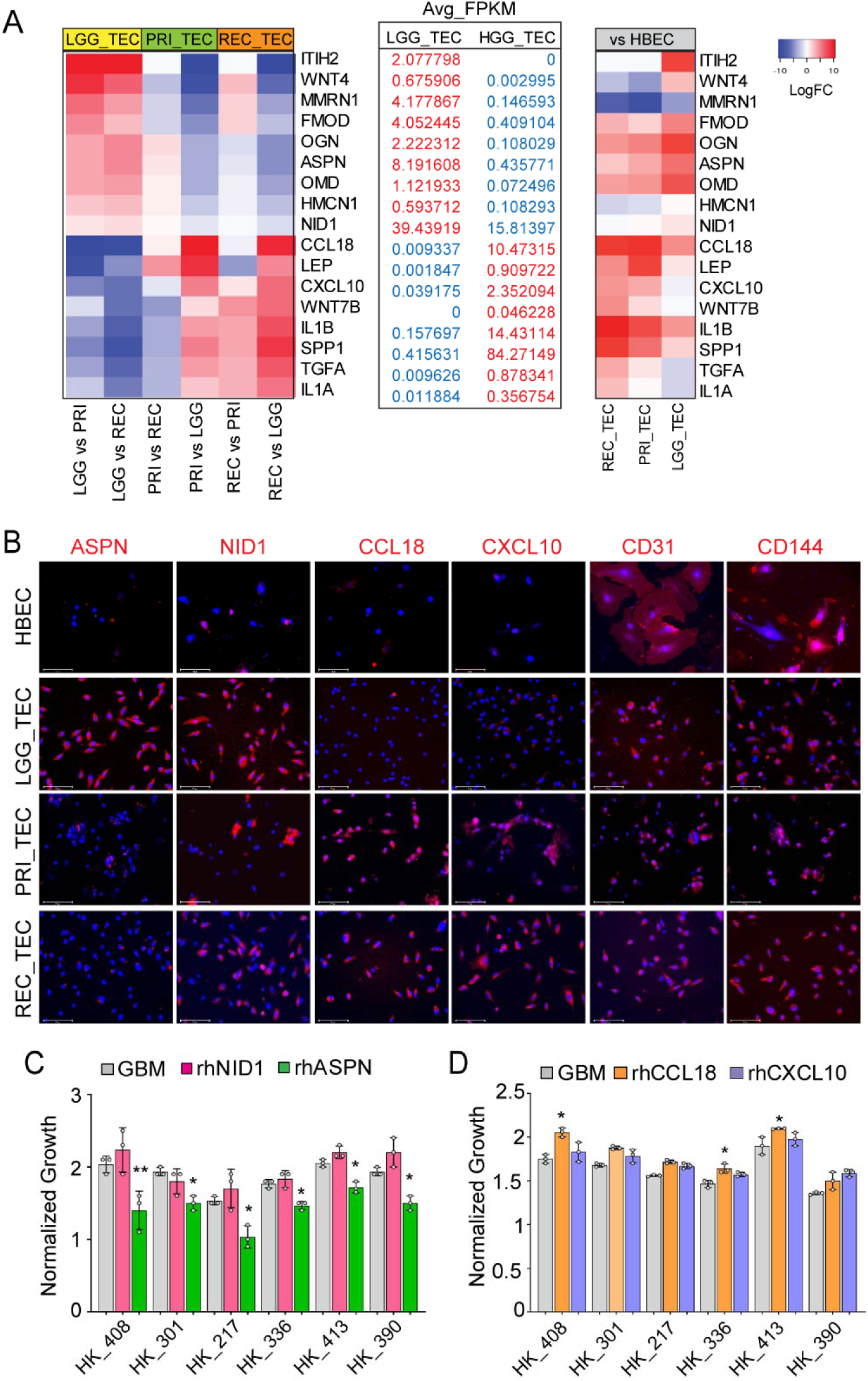
LGG-TEC and HGG-TEC show differential expression of cytokines, chemokines and proteoglycans. **A.** Heatmap of LogFC expression of significantly differentially expressed genes in LGG-TEC and HGG-TEC. Average FPKM values of each gene, and LogFC expression compared to HBEC. **B.** Immunostaining of LGG-TEC (ASPN, NID1) and HGG-TEC enriched genes (CCL18, CXCL10) and endothelial markers (CD31 and CD144/VE-CADHERIN) in cultured TEC and HBEC. Scale bars, 125μm. **C.** Normalized growth of wIDH GBM lines treated with recombinant NID1 and ASPN enriched in LGG-TEC. * p<0.05, ** p<0.005, one-way ANOVA **D.** Normalized growth of wIDH GBM lines treated with recombinant CCL18 and CXCL10 enriched in HGG-TEC. * p<0.05, ** p<0.005, one-way ANOVA

Further, we examined previously published bulk RNA sequencing data of freshly isolated CD31+ TEC from primary GBM tumors, and found that genes enriched in LGG-TEC were expressed at significantly low-or negligible levels in PRI-TEC compared to normal EC, and genes enriched in HGG-TEC were significantly upregulated in PRI-TEC corroborating the findings from cultured cells (Figure S4A)^8^. We also analyzed the expression of these genes in single-cell RNA-sequencing data of CD31+ TEC isolated from core and edge of primary GBM tumors. LGG-TEC-enriched genes (ITIH2, WNT4, FMOD, OGN, ASPN) showed very minimal expression in CD31+ TEC, whereas HGG-TEC-enriched genes especially IL1B and SPP1 were highly enriched in TEC from both core and edge of these primary GBM tumors (Figure S4B, C)^13^.

Next, we examined the expression of select candidates with high FPKM values in IVY_GAP database to determine the specific histological regions they were enriched in the tumors. LGG-TEC-enriched ASPN and NID1 were significantly higher in the microvascular proliferation regions compared to others, whereas HGG-TEC enriched chemokines CCL18 and CXCL10 did not exhibit significant enrichment in any specific region of the tumor (Figure S4D). To further validate this differential expression, we performed immunostaining on TEC cultures. As expected, ASPN and NID1 were highly expressed in LGG-TEC compared to HGG-TEC and HBEC. On the other hand, CCL18 and CXCL10 were expressed in HGG-TEC, but showed very low expression in LGG-TEC and HBEC, confirming the findings from RNA-sequencing (Figure 4B). Collectively, these data support the notion that LGG- and HGG-TEC are heterogeneous and show differential expression of genes including extracellular matrix proteins and cytokines.

We next tested whether these LGG- and HGG-TEC enriched genes differentially regulated the growth of GBM cells. Of the 4 candidates tested, ASPN enriched in LGG-TEC significantly inhibited the growth of tumor cells (N=6 wIDH GBM lines), whereas NID1 did not alter the growth of any of the tumor lines (Figure 4C). CCL18 enriched in HGG-TEC showed growth enhancing effect on two GBM lines but did not alter the growth of others, and CXCL10 did not affect the growth of tumor cells (Figure 4D). Based on these results, we hypothesized that the growth inhibitory effect of LGG-TEC on wIDH GBM cells is potentially mediated by ASPN.

### TEC express SLC1A1 transporter and uptake D-2HG

ASPN is an extracellular matrix protein that belongs to small-leucine rich proteoglycan (SLRP) family, and reported to play both tumor suppressive and oncogenic roles in different types of cancer^14, 15^. We first confirmed that ASPN is indeed enriched in LGG tumors by immunostaining tumor sections obtained from Grades II/III mIDH LGG (n=3) and Grade IV wIDH primary (n=3) and recurrent (n=3) GBM. Co-staining of ASPN with Tomato lectin to label blood vessels showed vascular expression of ASPN in LGG tumors, but minimal expression in HGG tumors (Figure 5A and Figure S5A).

**Figure 5:**
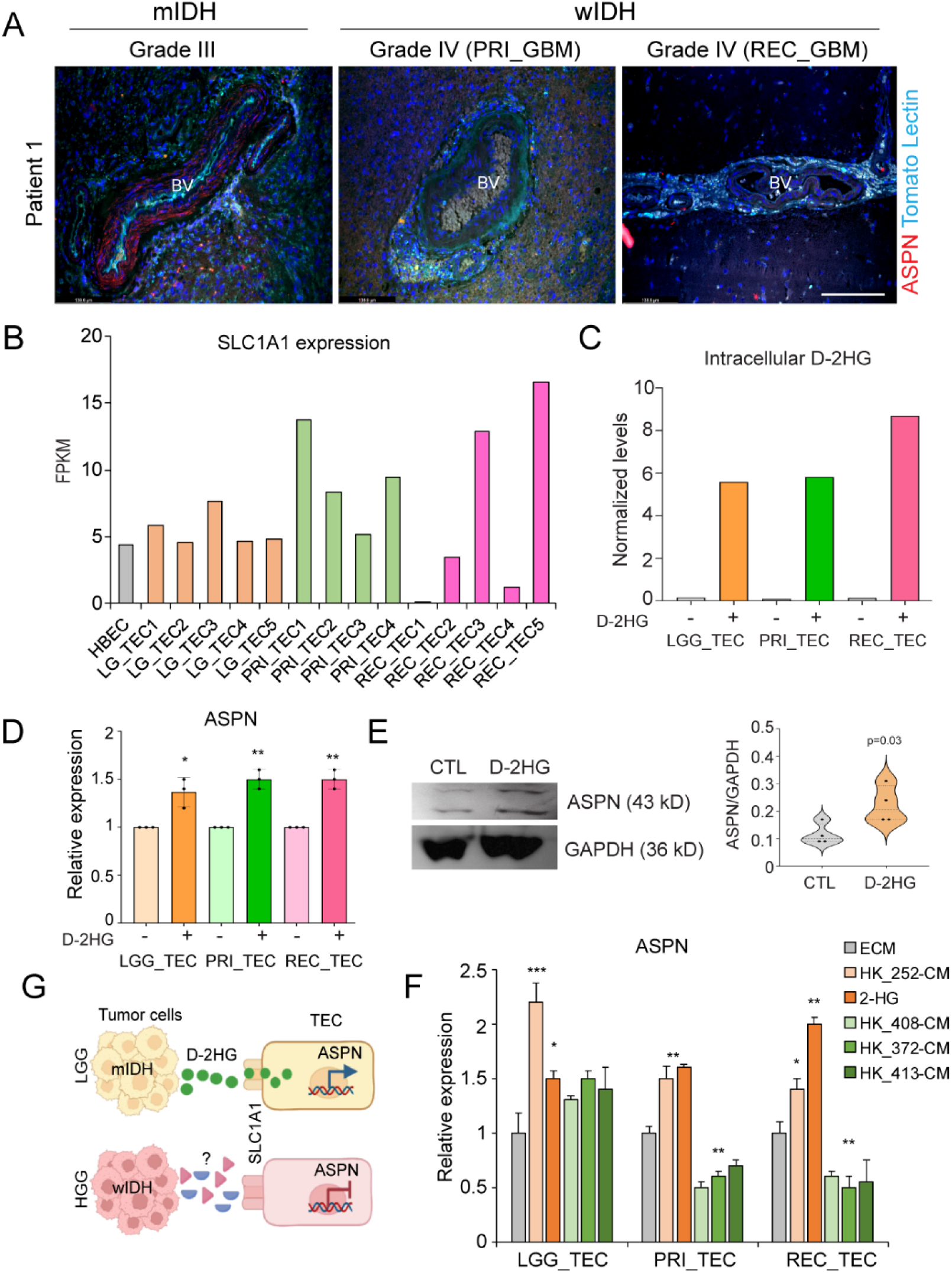
ASPN is highly expressed in vessels from mIDH LGG. **A.** Immunostaining of ASPN (red) and Tomato Lectin (blood vessels, cyan blue) in patient tumor tissue. Scale bars,150μm. **B.** FPKM expression of SLC1A1 in cultured HBEC and TEC **C.** Normalized intracellular levels of D-2HG in TEC cultured for 72 hours. **D.** Relative expression of ASPN in control and D-2HG-treated TEC. N=3 independent experiments, * p<0.05 and ** p<0.005, one-way ANOVA **E.** Immunoblot of ASPN and GAPDH in Control and D-2HG treated TEC. Quantitation of ASPN protein normalized to GAPDH. * p=0.03, unpaired t-test. **F.** Relative expression of ASPN in TEC treated with 2-HG, and conditioned media from mIDH astrocytoma (252) and wIDH (408, 372 and 413) GBM cells. **G.** Schematic illustrates the putative model for differential expression of ASPN between LGG and HGG-TEC

Since LGG-TEC cultures were all derived from mIDH tumors, we asked if ASPN expression is regulated by d-2-Hydroxyglutarate (2-HG), an oncometabolite secreted by mIDH tumor cells. A recent study reported that SLC1A1 is expressed by human umbilical vein endothelial cells (HUVEC) and facilitates the intracellular transport of 2-HG, which promotes endothelial migration and tumor angiogenesis^16^. We therefore examined our transcriptomic data to examine whether SLC1A1 is expressed by TEC. SLC1A1 was expressed by all TEC, whether freshly isolated from primary GBM tumors or cultured from LGG- and HGG-tumors, as well as by normal brain EC, albeit at varying levels (Figure 5B and Figure S5B). This suggested that TEC can uptake 2-HG from the tumor microenvironment via SLC1A1. We therefore treated our LGG-and HGG-TEC cultures with D-2HG that influxes into cells only in the presence of a transporter like SLC1A1. Strikingly, we observed high levels of intracellular D-2-HG in cell lysates from both LGG- and HGG-TEC indicating that TEC can indeed transport D-2-HG (Figure 5C).

### D-2HG promotes ASPN expression in TEC

Next, we tested whether treatment of TEC with D-2HG promoted ASPN expression. QRT-PCR analysis showed that D-2HG treatment increases ASPN expression in LGG- and HGG-TEC, which was further confirmed by immunostaining (Figure 5D and Figure S5C). We also performed immunoblotting of LGG-TEC treated with D-2HG and detected a significant increase in ASPN expression (Figure 5E). Additionally, we assessed whether D-2HG increases ASPN expression in a dose dependent manner. We found significant increase in ASPN expression with 20mM of D-2HG, and only a small increase at 10mM and 5mM indicating that there may be a dose dependent effect of D-2HG on ASPN expression (Figure S5D). Further, we also observed a significant increase in ASPN expression when LGG- and HGG-TEC were treated with conditioned media from a mIDH Grade IV astrocytoma tumor (252) line supporting the notion that 2-HG secreted by mIDH tumors promotes ASPN expression (Figure 5F). Since ASPN is expressed at relatively low levels in HGG-TEC from wIDH tumors, we wondered if GBM secreted factors inhibited ASPN expression. HGG-TEC cultured in conditioned media from 3 wIDH GBM lines showed significant reduction in ASPN expression, whereas LGG-TEC were unaffected indicating that ASPN expression in HGG-TEC is regulated by GBM cells (Figure 5F). Together, these results indicate that ASPN expression in TEC is differentially regulated between mIDH and wIDH tumors (Figure 5G).

### ASPN differentially inhibits the growth of wIDH GBM and mIDH astrocytoma tumors

Since ASPN is differentially regulated in wIDH and mIDH tumors, and exogenous addition of recombinant ASPN inhibited the growth of wIDH GBM lines, we asked whether it had opposing effects on the growth of wIDH and mIDH tumors. Indeed, ASPN expression significantly inhibited the growth of wIDH GBM tumor cells but had a small but significant growth promoting effect on mIDH astrocytoma cells (Figure 6A). To further confirm this, we measured EdU incorporation and found that wIDH GBM cells exposed to ASPN showed reduced proliferation whereas mIDH astrocytoma cells showed increased proliferation (Figure 6B and 6C).

**Figure 6:**
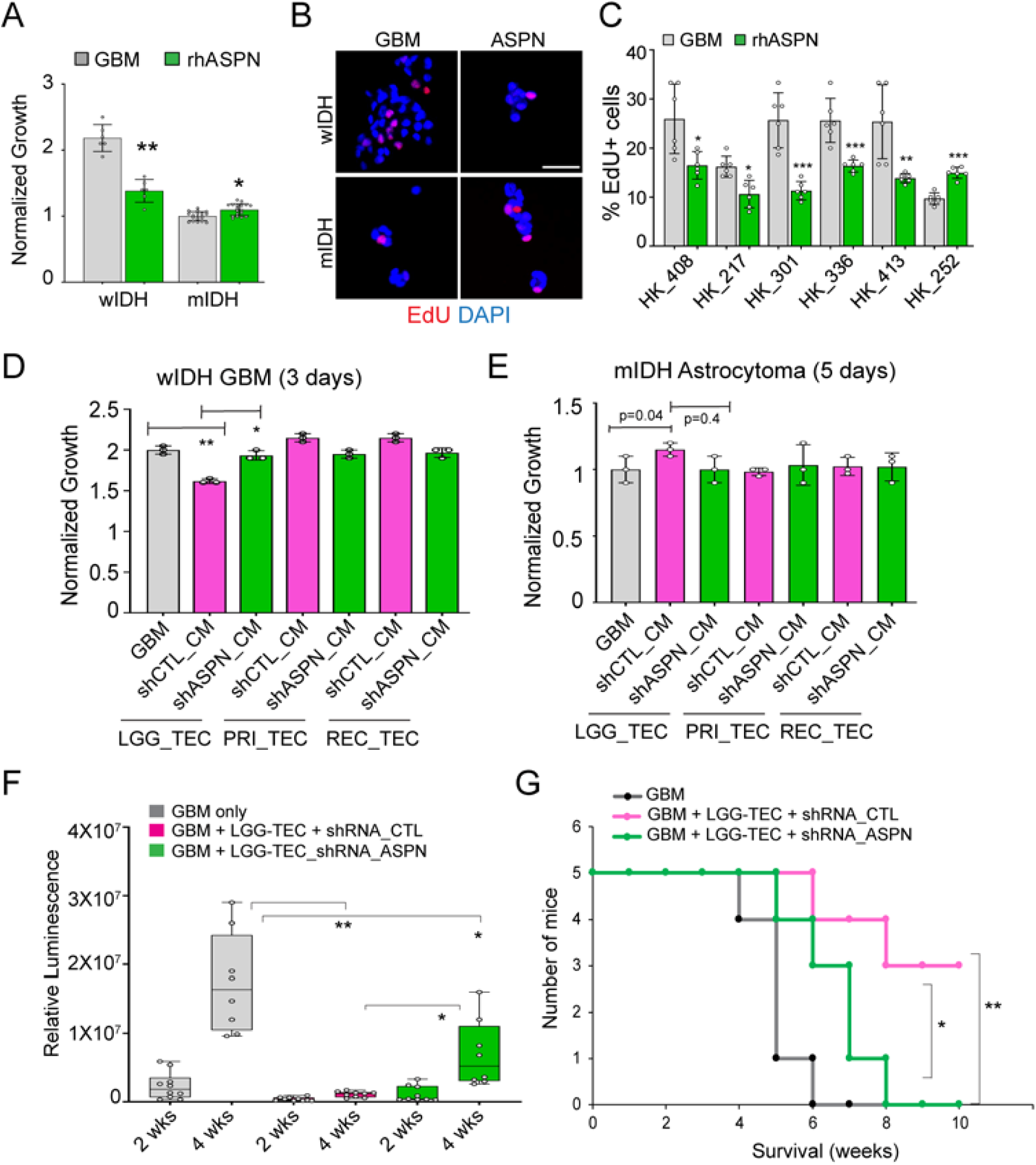
ASPN differentially controls the growth of mIDH astrocytoma and wIDH GBM. **A.** Normalized growth of wIDH GBM and mIDH astrocytoma cells treated with recombinant ASPN. *p<0.05, and ** p<0.005, unpaired t-test. **B.** Representative images of EdU (red) incorporation in wIDH GBM and mIDH astrocytoma cells treated with ASPN. Scale bars, 50μm. **C.** Quantitation of percentage of EdU+ cells in wIDH GBM (408, 217, 301, 336, 413) and mIDH (252) tumor cells. * p<0.05, ** p<0.005, and *** p<0.0005, unpaired t-test. **D.** Normalized growth of wIDH GBM cells treated with conditioned media from shRNA-CTL or shRNA-ASPN infected TEC. * p<0.05 and **p<0.005, one-way ANOVA. **E.** Normalized growth of mIDH tumor cells treated with conditioned media from shRNA-CTL or shRNA-ASPN infected TEC. P=0.04, one-way ANOVA. **F.** Box plots show relative luminescence from tumors in each condition at 2-and 4-weeks post-transplantation. N=5 mice per group. * p<0.05, ** p<0.005, one-way ANOVA and post-hoc t-test. **G.** Kaplan-Meier survival curve of mice co-transplanted with wIDH GBM and LGG-TEC infected with shRNA-CTL or shRNA-ASPN. * p<0.05, Log-rank test.

To functionally test if endogenous ASPN expressed by LGG-TEC is required for the growth-inhibitory effects on wIDH GBM cells, we used lentiviral shRNAs to knockdown ASPN. Knockdown (KD) efficiency was assessed by quantitative RT-PCR for ASPN mRNA and immunoblotting for ASPN protein (Figure S6A and S6B). We generated ASPN-KD and control (CTL) lines of both LGG- and HGG-TEC and verified by QRT-PCR (Figure S6C). Conditioned media (CM) from ASPN-KD cells partially rescued the growth inhibitory effect of LGG-TEC on wIDH GBM tumor cells (Figure 6D). However, ASPN-KD in HGG-TEC did not alter the growth of GBM cells. In addition, ASPN-KD in both LGG- and HGG-TEC did not significantly reduce the growth of mIDH astrocytoma cells (Fig 6E). These results suggested that LGG-TEC derived ASPN elicits a growth inhibitory effect specifically on wIDH GBM.

We next wanted to examine if ASPN inhibited growth of wIDH GBM tumors *in vivo.* As expected, co-transplantation of LGG-TEC with GBM cells significantly inhibited their growth, whereas co-injection of LGG-TEC lacking ASPN with GBM cells partially rescued the growth inhibitory effect corroborating the *in vitro* findings (Figure 6F). Survival analysis also showed that mice bearing tumors with LGG-TEC survived longer compared to GBM tumors only. However, mice bearing tumors with LGG-TEC lacking ASPN showed reduced survival indicating that ASPN in LGG-TEC is essential for the growth inhibitory effect on GBM tumors (Figure 6G). Collectively, these *in vitro* and *in vivo* findings strongly indicate that LGG-TEC derived ASPN inhibits growth of wIDH GBM tumors.

### ASPN inhibits wIDH GBM growth by modulating TGF***β***1 signaling

To determine the potential mechanism by which ASPN regulates wIDH GBM growth, we performed RNA-sequencing on ASPN-treated wIDH GBM (408) and mIDH astrocytoma (252) tumor cells. Differential expression analysis revealed a small number of genes regulated by ASPN in wIDH GBM, but a significantly greater number of genes in mIDH astrocytoma cells (Figure 7A). Of the top differentially expressed genes, most transcripts upregulated by ASPN in wIDH GBM were diminished in the mIDH tumor cells. Similarly, several transcripts downregulated by ASPN in wIDH GBM were either upregulated or showed no significant change in the mIDH tumor cells (Figure 7B). Gene ontology (GO) analysis showed that ASPN enriched for GPCR signaling, and downregulated TGFβ1 and ALK signaling in wIDH GBM cells, and conversely, upregulated these pathways in mIDH tumor cells (Figure 7C).

**Figure 7:**
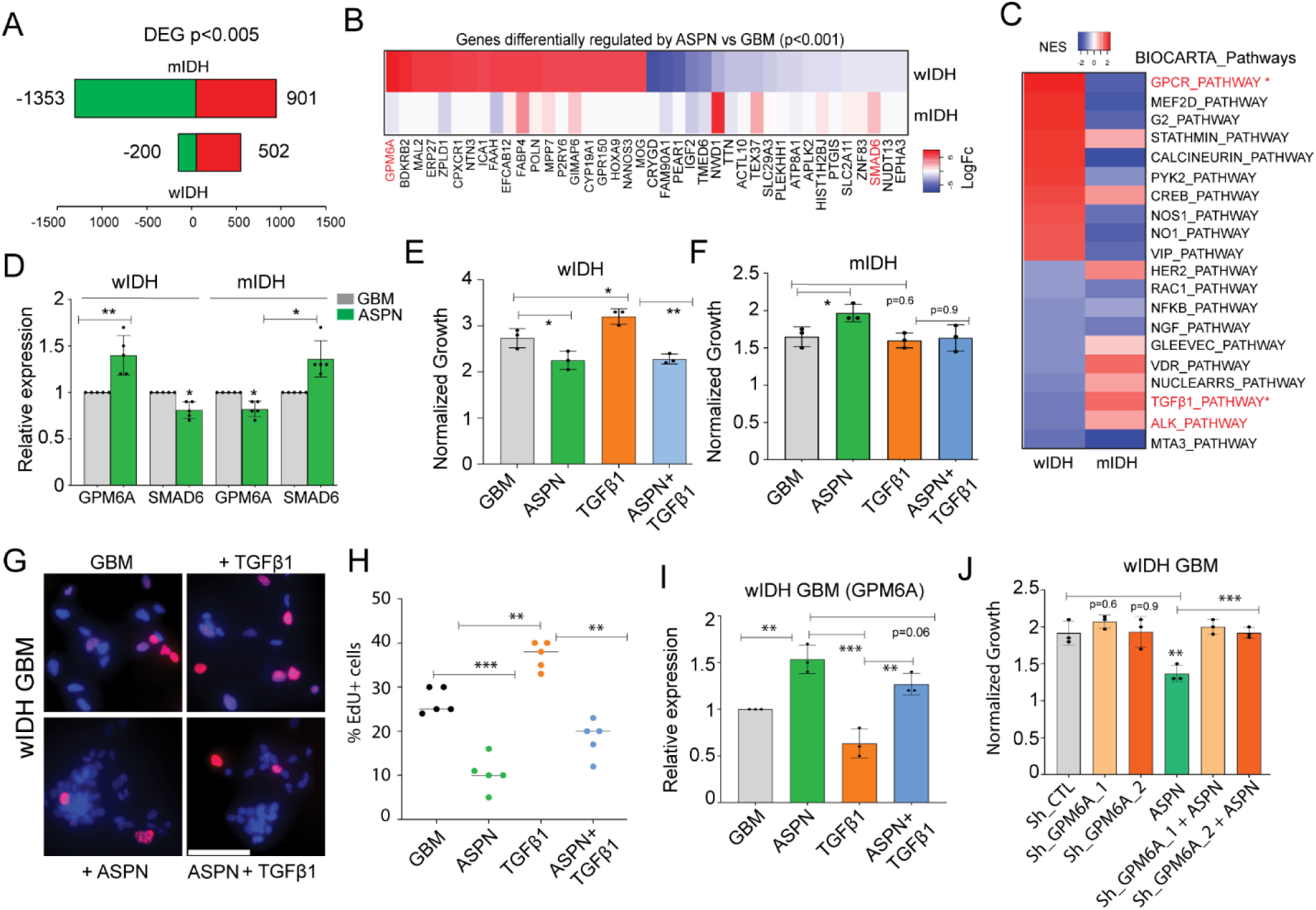
ASPN inhibits growth of wIDH GBM via TGF**β**1-GPM6A axis. **A.** Differentially expressed genes in wIDH GBM and mIDH astrocytoma treated with ASPN. **B.** Heatmap of ASPN-regulated genes between mIDH astrocytoma and wIDH GBM cells. **C.** Heatmap shows differentially regulated pathways between mIDH and wIDH tumor cells. **D.** Relative expression of GPM6A and SMAD6 in mIDH and wIDH tumor cells treated with ASPN. * p<0.05, and ** p<0.005, unpaired t-test. **E.** Normalized growth of wIDH cells treated with ASPN, TGFβ1 alone or in combination. * p<0.05, and ** p<0.005, one-way ANOVA **F.** Normalized growth of mIDH tumor cells treated with ASPN, TGFβ1 alone or in combination. * p<0.05, derived from one-way ANOVA. **G.** Representative images of EdU (red) incorporation in wIDH GBM cells treated with ASPN, TGFβ1 alone or in combination. Scale bars, 69.3μm. **H.** Quantitation of percentage of EdU+ cells in wIDH (408, 217, 301, 336, 413) and mIDH (252) tumor cells. ** p<0.005, and *** p<0.0005, one-way ANOVA. **I.** Relative expression of GPM6A in wIDH GBM cells treated with ASPN and TGFβ1 alone or in combination. ** p<0.005 and *** p<0.0005, one-way ANOVA. **J.** Normalized growth of shRNA-CTL and shRNA-ASPN infected wIDH GBM cells treated with ASPN. ** p<0.005, and *** p<0.0005, one-way ANOVA

ASPN has been previously reported to regulate TGFβ1 signaling^14, 17^. Consistent with this notion, we found that SMAD6, a TGFβ1 target gene was downregulated in wIDH GBM, and upregulated in mIDH tumor cells (Figure 7B). GPM6A, a highly enriched transcript in wIDH GBM upon ASPN treatment was previously reported to be suppressed by TGFβ1 signaling in mesothelial cells^18^. In addition, GPM6A is known to regulate MAPK signal transduction and recycling of GPCRs, and these pathways were increased with ASPN treatment in wIDH GBM (Figure 7C and Figure S7A)^19^. Based on these data, we hypothesized that ASPN inhibits the growth of wIDH GBM by modulating the TGFβ1-GPM6A axis.

To determine if TGFβ1 signaling regulates GBM growth, we treated wIDH GBM and mIDH astrocytoma cells with recombinant TGFβ1 either alone or in combination with recombinant ASPN. Expectedly, TGFβ1 treatment promoted growth, and addition of ASPN reversed this effect in wIDH GBM (Figure 7E). On the contrary, TGFβ1 did not have a significant effect on growth of mIDH astrocytoma cells (Figure 7F). We further confirmed this effect by measuring proliferation using EdU incorporation assay (Figure 7G, H). We also verified that TGFβ1 signaling was activated by immunostaining for pSMAD2/3 in wIDH and mIDH cells (Figure S7B). Moreover, we found that ASPN inhibited the expression of SMAD2/3 target genes including SMAD6, SMAD7 and ID1 downstream of TGFβ1 in wIDH GBM cells (Figure S7C). Together, these results indicated that ASPN antagonizes TGFβ1 signaling and its growth-promoting effect on wIDH GBM.

### Knockdown of GPM6A rescues the growth-inhibitory effect of ASPN in IDH1-wt GBM

To determine whether there is an inverse relationship between TGFβ1 and GPM6A downstream of ASPN, we measured the expression of GPM6A in TGFβ1- and ASPN-treated wIDH GBM cells. Expectedly, ASPN increased and TGFβ1 strongly inhibited the expression of GPM6A. The inhibitory effect of TGFβ1 on GPM6A expression was reversed by co-treatment with ASPN (Figure 7I). This supported our hypothesis that ASPN antagonizes TGFβ1 signaling to promote GPM6A expression and inhibit wIDH GBM growth.

Next, we asked whether we can rescue the growth-inhibitory effect of ASPN by blocking GPM6A expression. GPM6A knockdown in wIDH GBM and mIDH tumor cells was performed using shRNA constructs. Knockdown efficiency was validated by QRT-PCR and immunostaining (Figure S7D and S7E). Interestingly, GPM6A-KD alone did not have significant effects on growth of either wIDH or mIDH tumor cells. However, GPM6A-KD rescued the growth inhibition of ASPN in wIDH GBM but had no effect in mIDH tumor cells (Figure 7J and Figure S7F). These results indicate that GPM6A is essential for ASPN-mediated suppression of wIDH GBM growth. Collectively, our findings indicate that low-grade TEC-derived ASPN inhibits the growth and proliferation of wIDH GBM cells by regulating the TGFβ1-GPM6A axis (Figure 8).

**Figure 8:**
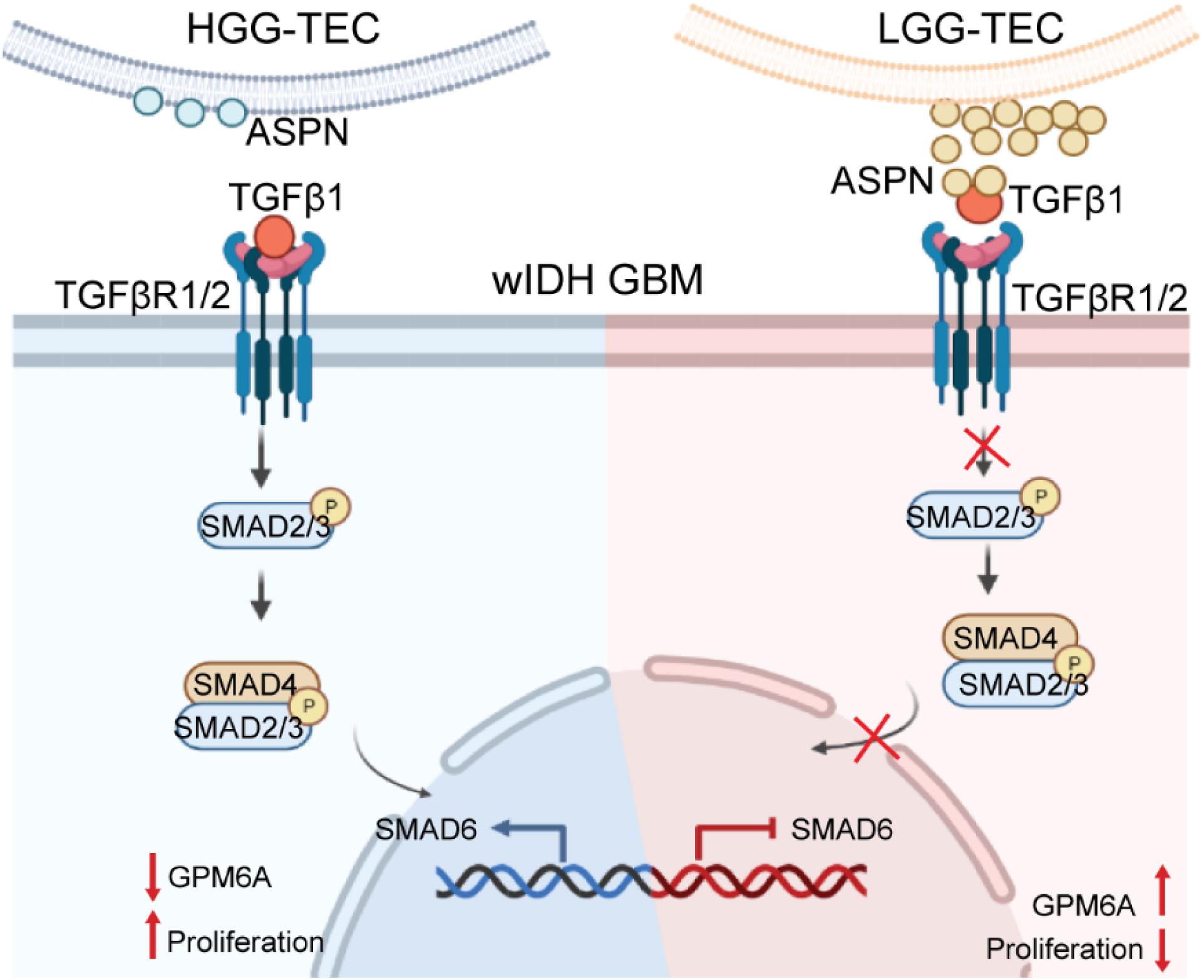
Model of differential regulation of TGFβ1 signaling and growth of wIDH GBM by LGG- and HGG-TEC.

## Discussion

Early transcriptomic profiling studies reported that TEC from LGG and HGG exhibit significant phenotypic and molecular heterogeneity from normal brain EC^5, 6, 11^. More recent single-cell RNA sequencing studies demonstrated the intratumoral heterogeneity of TEC derived from core and edge of primary GBM tumors, as well as from transdifferentiated ECs^7, 13^. These studies, while being a valuable resource, have not yielded insights into specific mechanisms by which TEC heterogeneity contributes to tumor growth and resistance.

In this study, we established TEC cultures from Grade II/III mIDH LGG and Grade IV wIDH HGG tumors to not only elucidate their molecular heterogeneity but also understand how this heterogeneity influences tumor growth and progression. By performing extensive transcriptomic sequencing of these cultured LGG- and HGG-TEC and human brain endothelial cells, we identified key molecular differences in their expression of extracellular matrix proteins, growth factors and cytokines. We also demonstrated that a few of these differentially expressed factors have distinct effects on the growth and proliferation of GBM cells derived from wIDH and mIDH tumors. This indicated that the mechanisms by which TEC control tumor growth may be different in LGG vs HGG and also dependent on the mutational status of the tumors as previously indicated^11^.

LGG- and HGG-TEC exhibit significant functional differences in their response to anti-angiogenic drugs, Bevacizumab (BVZ) and Sunitinib. In line with prior findings, HGG-TEC are resistance to both anti-angiogenic treatments, whereas LGG-TEC are sensitive to these drugs^20^. While these therapies have failed in clinic in improving patient survival outcomes, as they were predominantly tested on recurrent GBM patients, they may still hold some promise for LGG, and warrants further investigation.

Prior research indicated that radiation therapy disrupts the vasculature and exerts a broad range of effects including endothelial senescence, increased inflammation, immune cell recruitment and re-vascularization of the tumor^21, 22^. Here, we found that both LGG- and HGG-TEC are resistant to high-doses of radiation and proliferate similar to non-radiated cells. However, our analysis was limited to assessing proliferation for only a short duration of 3 days, and we did not assess the extent of DNA damage in TEC. Further experiments are needed to elucidate whether TEC are refractory to radiation in long-term culture or undergo senescence and display adaptive resistance. In addition, it remains undetermined how anti-angiogenic, radiation or chemotherapy alter the molecular landscape of TEC and in turn influence tumor growth.

A major and unexpected finding of this study is that LGG-TEC derived factors have differential effects on the growth of wIDH and mIDH tumors. While we validated the growth inhibitory effect of LGG-TEC on wIDH tumors *in vivo* in co-transplantation studies, we were not successful in growing mIDH tumors in orthotopic xenograft models. There is an unmet need in GBM research to develop methods to effectively transplant and grow LGG and HGG mIDH tumors *in vivo*.

Our differential gene expression analysis revealed ASPN as highly enriched in the mIDH LGG- TEC. ASPN expression is dysregulated in several cancers, and has been reported to act as an oncogene in pancreatic, colorectal, gastric and prostate cancer, and as a tumor suppressor in triple negative breast cancer^23^. Moreover, it regulates several signaling pathways including TGFB, EGFR and CD44 pathways to control tumor proliferation, migration and invasion^23^. The role of ASPN has not been previously described in GBM or its expression in glioma vasculature. Our data demonstrated that ASPN is highly enriched in tumor vessels of LGG compared to HGG, and its expression is regulated by 2-hydroxyglutarate (2-HG), an oncometabolite secreted by mIDH tumors. More interestingly, ASPN expression in HGG-TEC was suppressed by treatment with conditioned media of wIDH tumor cells, but not in LGG-TEC. This indicates that

ASPN acts as a tumor suppressor, and its expression is differentially regulated in TEC from mIDH LGG and wIDH HGG tumors. The specific mechanism by which ASPN expression is suppressed in HGG-TEC, and what other functions it may serve in GBM biology remains to be determined. We speculate that inhibition of ASPN expression, though is a prerequisite to the establishment or progression of GBM.

ASPN treatment differentially altered the transcriptional landscape of wIDH and mIDH tumors. Several genes upregulated by ASPN in wIDH GBM cells are diminished in expression in mIDH tumor cells including TGFβ1 and GPCR pathway-associated genes. Glycoprotein M6A (GPM6A), the most significantly upregulated gene in wIDH GBM cells, is markedly downregulated in mIDH tumor cells upon ASPN treatment. On the other hand, SMAD6, a downstream target of TGFβ1 signaling is downregulated by ASPN in wIDH GBM, but enhanced in mIDH tumor cells indicating that ASPN differentially influences these signaling pathways. TGFβ1 was previously reported to modulate the expression of GPM6A in mesothelial cells of the liver^24^. In line with this, our findings also show that TGFβ1 treatment reduces GPM6A expression, whereas ASPN increases GPM6A by blocking TGFβ1 signaling in wIDH GBM cells. Furthermore, GPM6A knockdown rescues the growth-inhibitory effect of ASPN in wIDH GBM cells, but has no effect on mIDH tumor cells suggesting that they all function in a single axis to control growth of wIDH GBM cells. One potential advantage of downregulating ASPN in HGG- TEC by wIDH GBM cells could be that it suppresses TGFβ1 signaling, a known effector signaling molecule of immunosuppression that aids in tumor cell escape from immune surveillance and promotes tumor progression^25^. Future studies will be needed to investigate whether ASPN overexpression blocks TGFβ1 signaling in GBM tumors.

In conclusion, our study revealed the molecular and functional heterogeneity between LGG- and HGG-TEC, and identified ASPN expressed by LGG-TEC as a potential regulator of TGFβ1 signaling-mediated tumor growth.

## Acknowledgements

The authors thank the UCLA BTTR, JCCC Flow Cytometry Core, UNGC, and the TCGB core for technical contributions.

## Authorship

S.D.M and H.I.K conceptualized the study. S.D.M, H.Q., L.E., D.W., A.P., A.G.A., T.L. performed experiments. F.G., R.K. analyzed the sequencing data and performed bioinformatics analyses. A.L. provided reagents. S.D.M and H.I.K wrote and edited the manuscript. All authors contributed to the revision of the manuscript.

## Data availability

All sequencing data has been submitted to Gene Expression Omnibus, and are available with the Accession number GSE236571.

