## Supplemental File for "Low- and high-grade glioma endothelial cells differentially regulate tumor growth"

**Supplemental Figures**

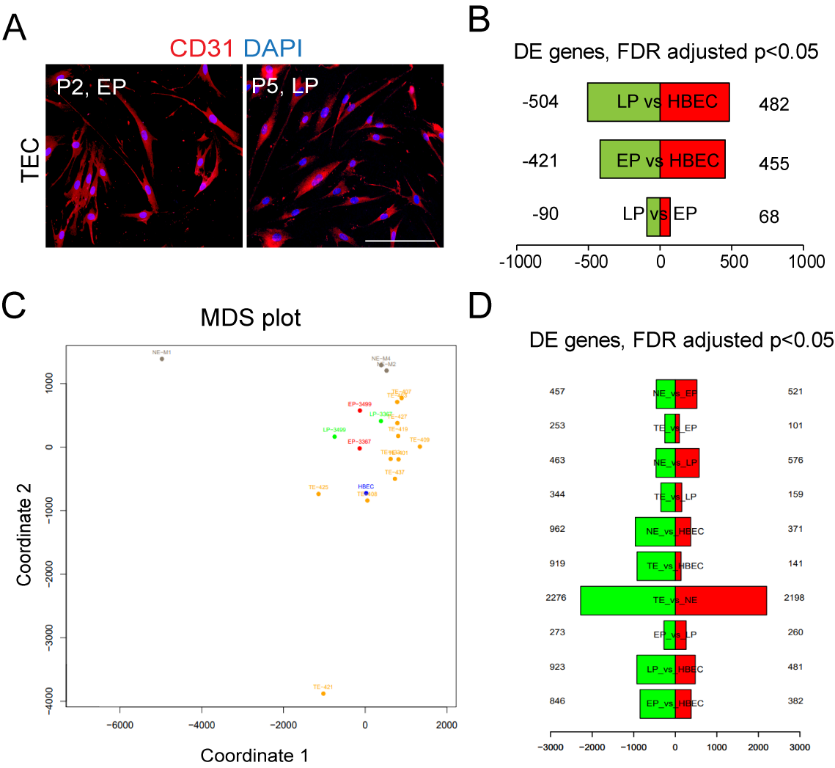

**Figure S1: Characterization of cultured TEC**

A. Immunostaining of CD31 (red) in cultured tumor endothelial cells (TEC) at early (EP) and late (LP) passage. Scale bars,150μm.
B. Differentially expressed genes between HBEC, early and late passage TEC.
C. MDS plot of cultured early (EP) and late passage (LP) TEC and freshly isolated tumor endothelial cells (TE) from primary GBM tumors and normal brain EC (NE).
D. Differentially expression genes between NE, TE, HBEC and early (EP) and late (LP) passage TEC.

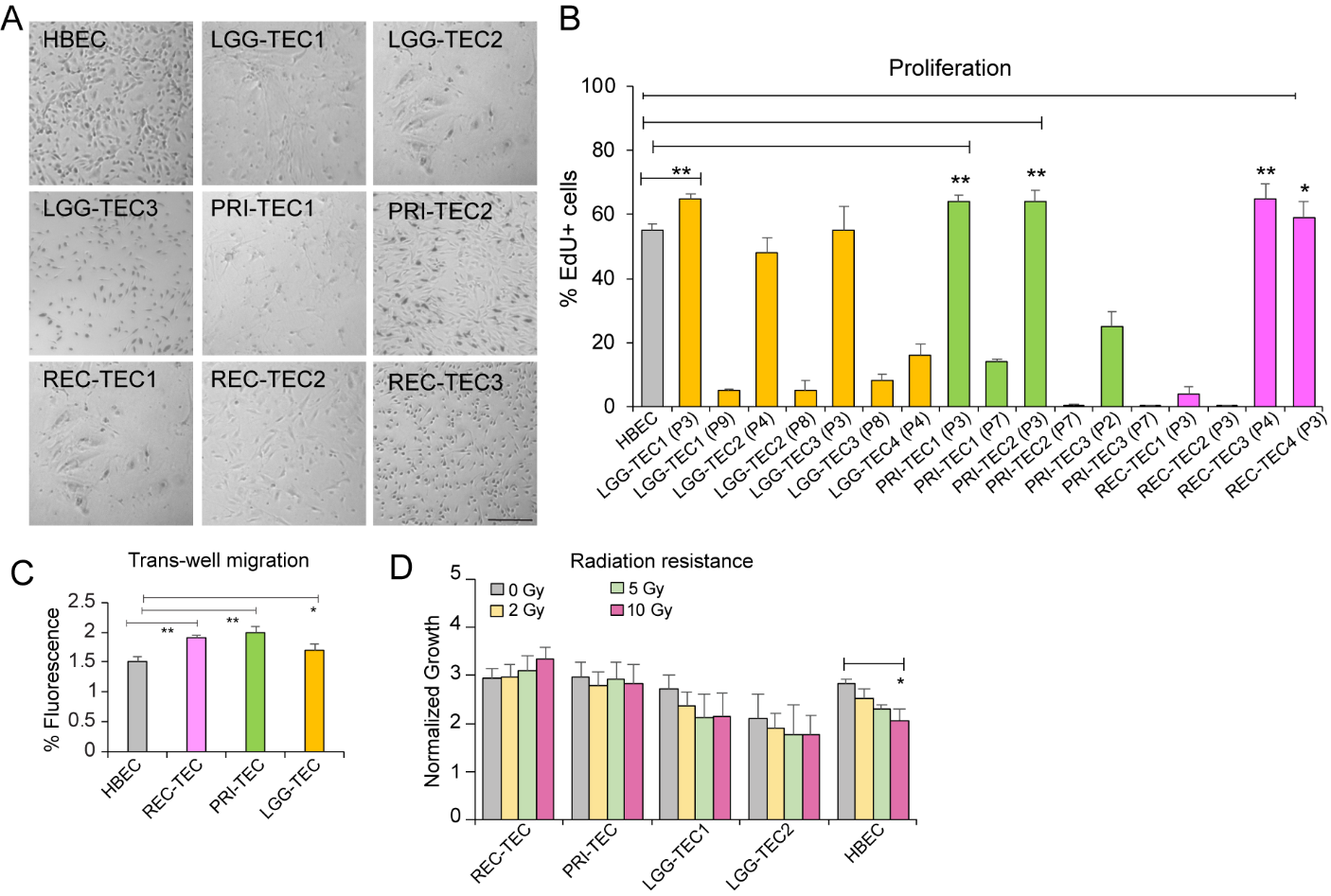

**Figure S2: Functional differences between LGG- and HGG-TEC**

A. Images of HBEC and cultured LGG-, PRI- and REC-TEC. Scale bars,150μm.
B. Quantitation of percentage of EdU incorporation in cultured HBEC, LGG-, PRI- and REC-TEC lines. * p<0.05, one-way ANOVA.
C. Quantitation of fluorescence indicating migration distance of cultured TEC. * and ** p<0.05, p<0.005, one-way ANOVA.
D. Normalized growth and proliferation of of HBEC and TEC in response to various doses of radiation. * p<0.05, one-way ANOVA.

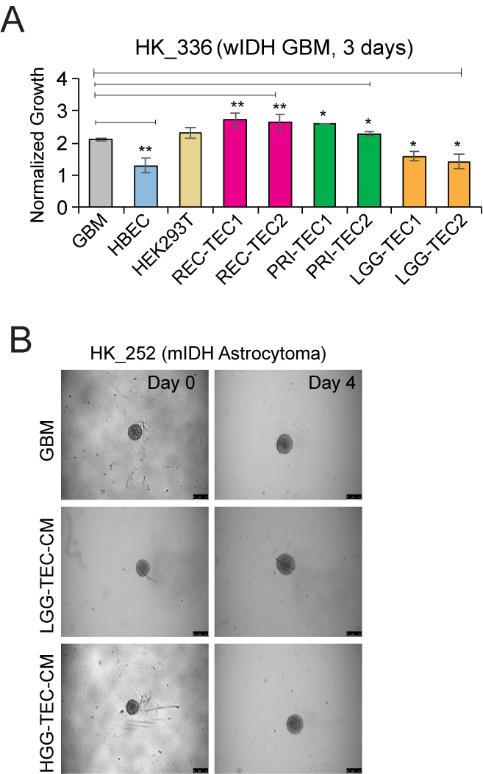

**Figure S3: LGG- and HGG-TEC differentially regulate the growth of mIDH astrocytoma and wIDH tumor cells**

A. Normalized growth of wIDH GBM cells in conditioned media of cultured TEC. * and ** p<0.05, p<0.005, one-way ANOVA.
B. Images of mIDH astrocytoma spheroids cultured in conditioned media of LGG- and HGG-TEC. Scale bars, 250μm.

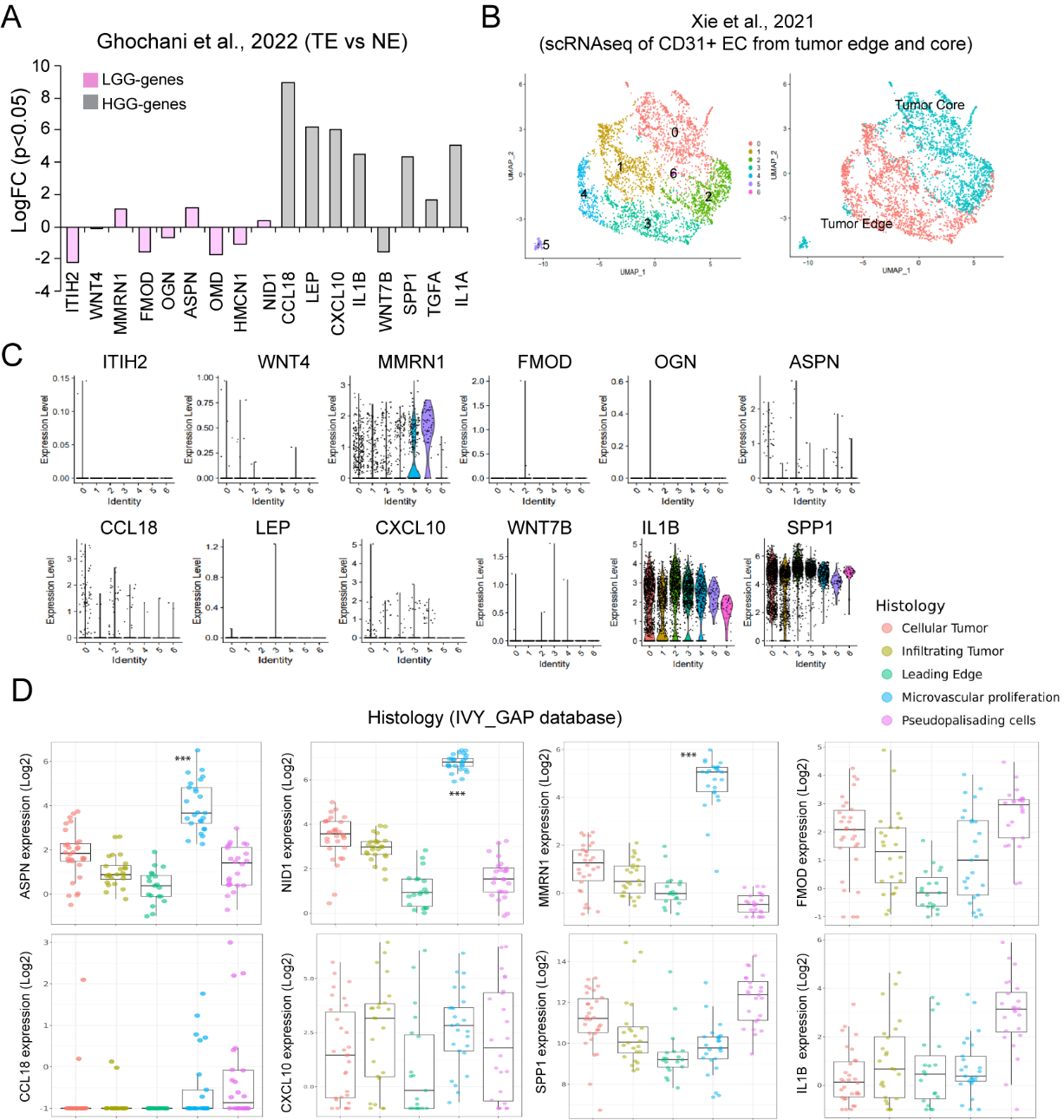

**Figure S4: LGG- and HGG-TEC shows differential expression of growth factors, cytokines and extracellular matrix proteoglycans.**

A. LogFC values of cultured LGG- and HGG-TEC specific genes in freshly isolated primary GBM tumor endothelial cells (TEC) compared to normal brain EC (NE)
B. UMAP plot of CD31+ EC clusters derived from core and edge of primary GBM tumors
C. Expression of LGG- and HGG-TEC genes in clusters from the scRNAseq data
D. Log2 expression values of LGG- and HGG-genes in different histological regions of tumors in IVY_GAP database. *** indicates p<0.0005, pairwise t-test with Bonferroni correction.

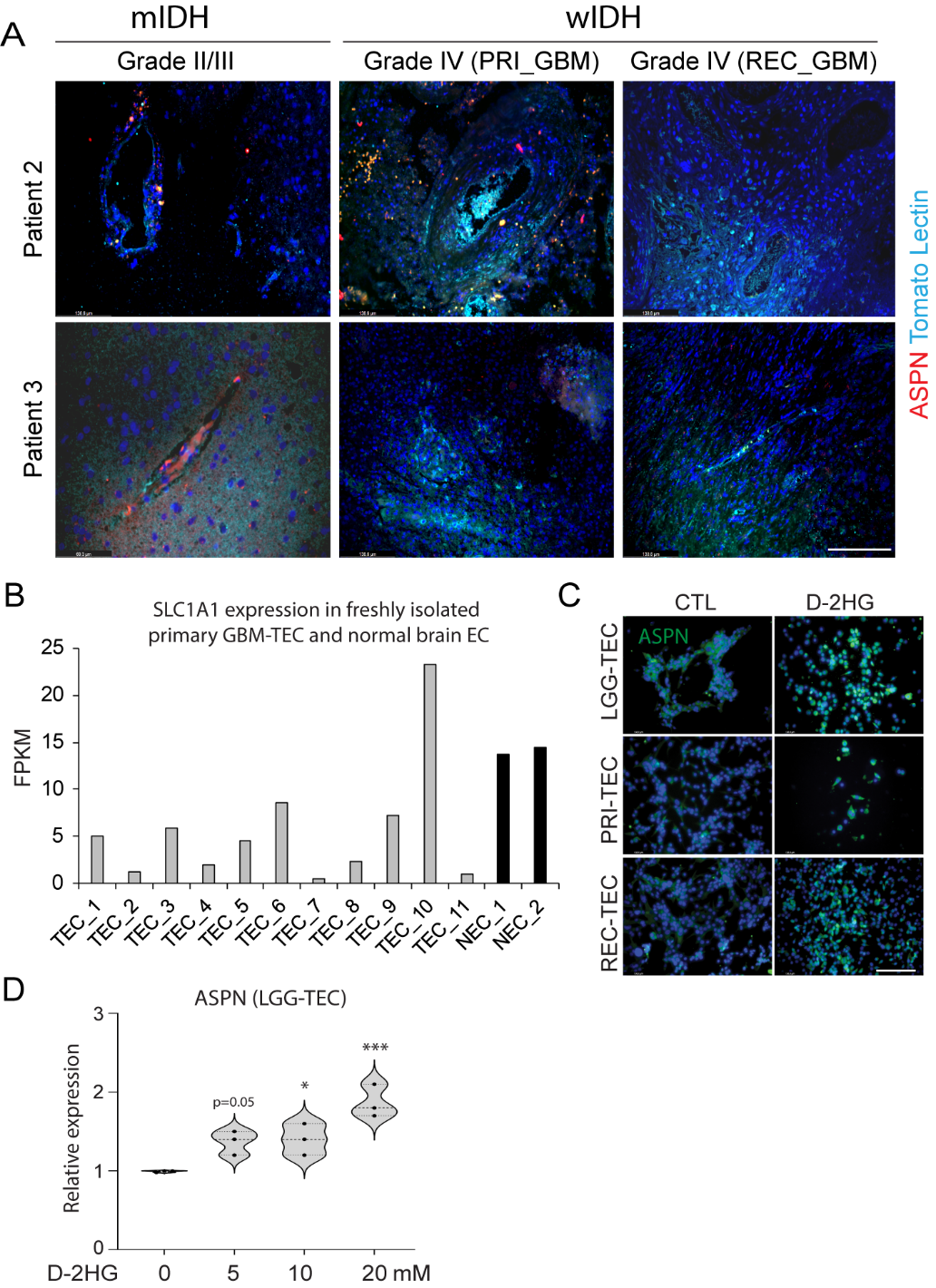

**Figure S5: ASPN expression is enriched in IDH mutant LGG and regulated by D-2HG**

A. Immunostaining of ASPN (red) and Tomato Lectin (Cyan blue, blood vessels) in mIDH LGG and wIDH primary and recurrent GBM tumor tissue. Scale bars,150μm.
B. FPKM values of SLC1A1 in freshly isolated TEC from primary GBM and normal brain EC.
C. Immunostaining of ASPN in cultured TEC treated with D-2HG. Scale bars,150μm.
D. Relative expression of ASPN in LGG-TEC1 treated with different concentrations of D-2HG. * and ** indicates p<0.05, P<0.005, one-way ANOVA.

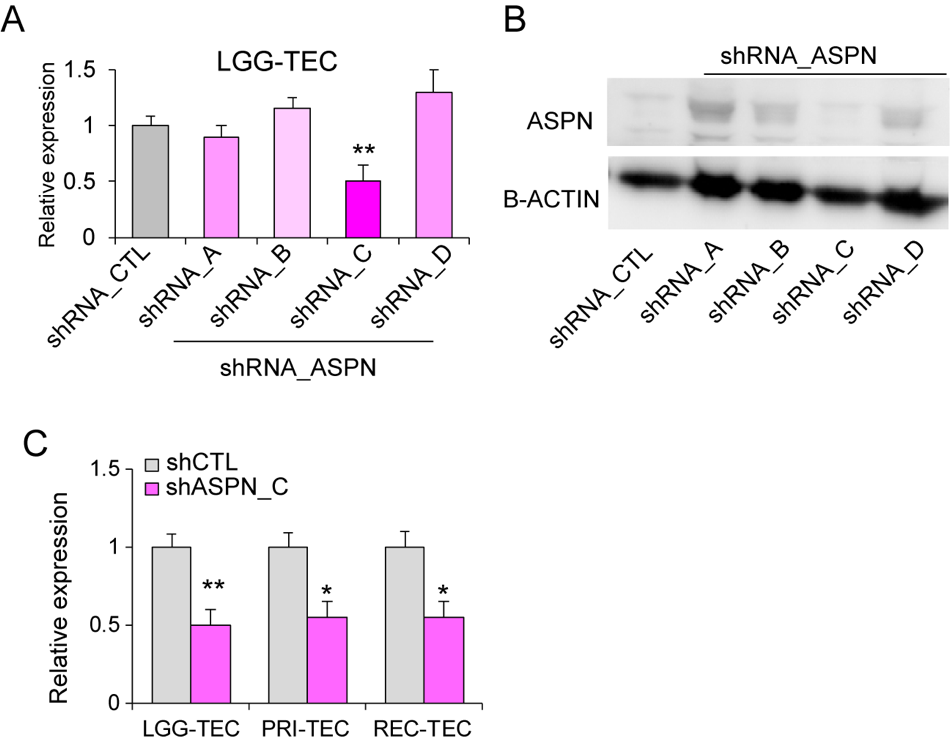

**Figure S6: ASPN knockdown in cultured TEC**

A. Relative expression of ASPN in cultured LGG-TEC infected with CTL and ASPN shRNAs. *p<0.05, one-way ANOVA
B. Immunoblot of ASPN and B-ACTIN in cultured LGG-TEC infected with CTL and ASPN shRNAs.
C. Relative expression of ASPN in cultured LGG-TEC, PRI-TEC and REC-TEC. *p<0.05, **p<0.005, unpaired t-test.

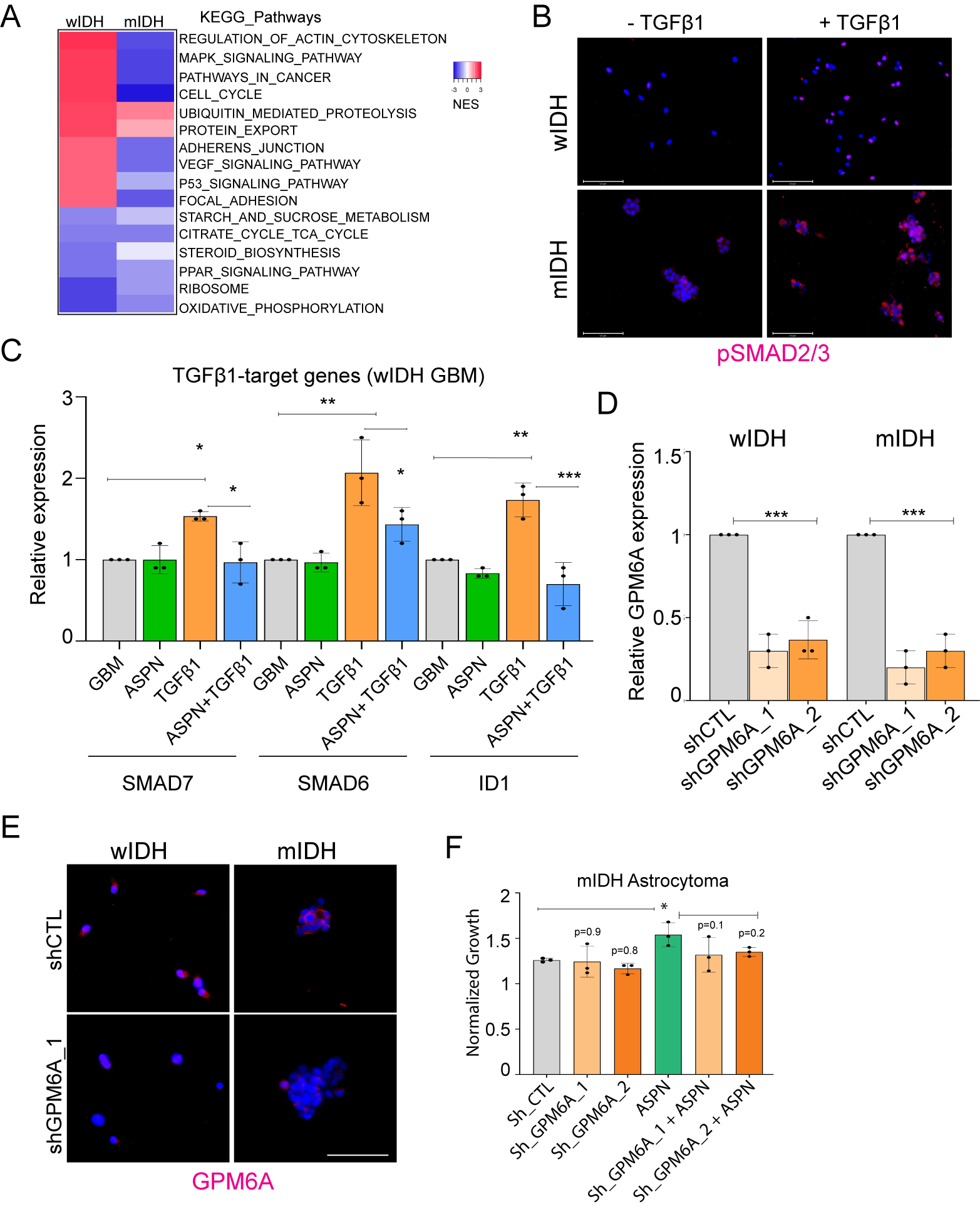

**Figure S7: ASPN inhibits wIDH GBM growth via modulation of TGFΒ1-GPM6A signaling**

A. Heatmap shows normalized enrichment scores (NES) of KEGG_pathways in mIDH astrocytoma and wIDH GBM treated with ASPN.
B. Immunostaining of phospho-SMAD2/3 (red) in mIDH astrocytoma and wIDH GBM cells treated with recombinant TGFβ1. Scale bars, 125μm
C. Relative expression of TGFβ1-target genes in wIDH GBM cells treated with ASPN, TGFβ1 alone or in combination. * p<0.05, ** p<0.005, *** p<0.0005, one-way ANOVA.
D. Relative expression of GPM6A in mIDH astrocytoma and wIDH GBM cells infected with CTL and GPM6A shRNAs. *** p<0.0005, one-way ANOVA.
E. Immunostaining of GPM6A in GPM6A in mIDH astrocytoma and wIDH GBM cells infected with CTL and GPM6A shRNA. Scale bars, 125μm
F. Normalized growth of control and GPM6A-knockdown mIDH tumor cells treated with ASPN. P-values derived from one-way ANOVA, * p<0.05.

**Supplemental Tables**

**Table S1: Patient samples**

| **TEC ID** | **Glioma Classification** | **Glioma Grade** |
| --- | --- | --- |
| LGG-TEC1 | Oligodendroglioma | Grade II, IDH mutant, 1p 19q deletion |
| PRI-TEC1 (EP/LP1) | Glioblastoma | Grade IV, IDH wildtype |
| REC-TEC1 (EP/LP2) | Glioblastoma | Grade IV, IDH wildtype |
| PRI-TEC2 | Glioblastoma | Grade IV, EGFR amplified |
| LGG-TEC2 | Oligodendroglioma | Grade III, IDH mutant |
| LGG-TEC3 | Astrocytoma | Grade III, IDH mutant |
| LGG-TEC4 | Glioma | Grade II, IDH mutant |
| LGG-TEC5 | Astrocytoma | Grade III, IDH mutant |
| LGG-TEC6 | Glioma | Grade II, IDH mutant |
| PRI-TEC3 | Glioblastoma | Grade IV, IDH wildtype |
| PRI-TEC4 | Glioblastoma | Grade IV, IDH wildtype |
| PRI-TEC5 | Glioblastoma | Grade IV, IDH wildtype |
| PRI-TEC6 | Glioblastoma | Grade IV, IDH wildtype, EGFR amplified |
| PRI-TEC7 | Glioblastoma | Grade IV, IDH wildtype |
| REC-TEC2 | Glioblastoma | Grade IV, IDH wildtype, EGFR amplified |
| REC-TEC3 | Glioblastoma | Grade IV, IDH wildtype |
| REC-TEC4 | Glioblastoma | Grade IV, IDH wildtype |
| REC-TEC5 | Glioblastoma | Grade IV, IDH wildtype |

**Table S2: Human Oligonucleotides**

| **Genes** | **Forward primer** | **Reverse primer** |
| --- | --- | --- |
| **18srRNA** | GGCCCTGTAATTGGAATGAGTC | CCAAGATCCAACTACGAGCTT |
| **PECAM1/CD31** | CCAAGGTGGGATCGTGAGG | TCGGAAGGATAAAACGCGGTC |
| **CDH5** | AAGCGTGAGTCGCAAGAATG | TCTCCAGGTTTTCGCCAGTG |
| **VWF** | CCGATGCAGCCTTTTCGGA | TCCCCAAGATACACGGAGAGG |
| **FLT1** | GAAAACGCATAATCTGGGACAGT | GCGTGGTGTGCTTATTTGGA |
| **ENG** | CGCCAACCACAACATGCAG | GCTCCACGAAGGATGCCAC |
| **CD248** | TGGTGCCAACGTGTGTCTTTT | AGCGATAGCAGTCAGTGATGC |
| **MCAM** | AGCTCCGCGTCTACAAAGC | CTACACAGGTAGCGACCTCC |
| **DES** | GAGACCATCGCGGCTAAGAAC | GTGTAGGACTGGATCTGGTGT |
| **ACTA2** | CTATGAGGGCTATGCCTTGCC | GCTCAGCAGTAGTAACGAAGGA |
| **PDGFRB** | AGCACCTTCGTTCTGACCTG | TATTCTCCCGTGTCTAGCCCA |
| **ASPN** | CTCTGCCAAACCCTTCTTTAGC | CGTGAATAGCACTGACATCCAA |
| **ID1** | CTGCTCTACGACATGAACGG | GAAGGTCCCTGATGTAGTCGAT |
| **SMAD6** | CCTCCCTACTCTCGGCTGTC | GGTAGCCTCCGTTTCAGTGTA |
| **SMAD7** | GGACGCTGTTGGTACACAAG | GCTGCATAAACTCGTGGTCATTG |
| **GPM6A** | ATTCCCTATGCCTCTCTGATTGC | GCCATCTCAAAGTAGGTTTGCAG |

**Supplementary methods:**

1. **Isolation of tumor and endothelial cells from patient tissues:** Human brain tumor tissue samples were obtained from surgical resections from patients under an approved University of California, Los Angeles (UCLA) Institutional Review Board (IRB) protocol. Information on patient diagnoses can be found in Tables S1A. Single cell suspension of the tissue (0.3-0.5g) was obtained by mincing into smaller pieces, followed by enzymatic digestion in Collagenase II and Collagenase IV in Hibernate A media at 37⁰C for 30 min. Following enzymatic digestion, the cell suspension was passed through 100 and 70µm filters and cells were pelleted at 1000xg for 5mins. Cellular debris and red blood cells (RBC) was removed by pelleting the cell suspension in 1.5ml of DMEM/F12, and 1.5ml of 1X Percoll solution. RBCs were pelleted by centrifugation of this suspension at 1000xg for 5min. 1.5ml of 4X percoll buffer was gradually added to the remaining supernatant to facilitate a shift in osmolality that would selectively allow live cells to pellet, and centrifuged at 3000xg for 7min. Cells were gradually re-introduced to normal salt concentrations by addition of 10ml of DMEM:F12, and passed through a 40µm filter, and pelleted by centrifugation at 1000xg for 5min. The cells were then re-suspended in 1ml of phosphate buffered saline (PBS) containing 0.1% Bovine Serum Albumin and the number of cells and their viability were assessed. **MACS sorting of CD31+ TEC:** Cell suspension was resuspended in CD31-antibody conjugated Dynabeads in 1ml of PBS with 0.1 % BSA and incubated for 20 minutes. CD31+ TEC were enriched and sorted according to the manufacturer’s protocol. Remaining CD31-negative fraction containing tumor cells were pelted by centrifugation at 1000g for 5mins. Cells were counted and viability was assessed by Tryphan blue staining and plated in endothelial growth media (ECM) or GBM media for expansion.
2. **RNA sequencing and analysis:** RNA extraction from gliomaspheres and tumor endothelial cell lines was performed using Qiagen RNeasy microkit. RNA quality was assessed using Bioanalyzer and only samples with a RIN score >8.0 were sequenced. RNA samples were pooled and barcoded, and libraries were prepared using TruSeq Stranded RNA (100ng) + Ribozero Gold. Paired-end 150bp reads were aligned to the latest human_hg38 reference genome using the STAR spliced read aligner. Total counts of read-fragments aligned to known gene regions within the human hg38 refSeq reference annotation was used as the basis for quantiﬁcation of gene expression. Diﬀerentially expressed genes were identiﬁed using DESeq and edgeR, and ranked based on False Discovery Rate (FDR Bejamini Hochberg adjusted p-values of ≤ 0.01). Gene Set Enrichment Analysis (GSEA) was carried out to determine the gene signatures differentially regulated between groups and represented as heatmaps.
3. **Quantitative RT-PCR:** RNA was isolated with RNeasy micro kit (QIAGEN), and 100-500g was used for first-strand cDNA synthesis using random primers and Superscript Reverse Transcriptase (Invitrogen). qRT–PCR was performed using Power SYBR Green PCR Master Mix (Applied Biosystems). The relative expression of genes was normalized using 18srRNA as the housekeeping gene. All experiments were repeated at least 3 times, and data is represented as mean ± SD. Primers are listed in the Table S1B.
4. **Immunofluorescence staining:** 5μm FFPE GBM patient tissue sections were incubated with primary antibodies overnight at 4°C after deparaffinization, rehydration, antigen retrieval and blocking in PBS with 2% BSA. Sections were then incubated with species-appropriate goat/donkey secondary antibodies coupled to AlexaFluor dyes (488, 568 or 647, Invitrogen) and Hoechst dye for nuclear staining for 2 hrs at RT. VECTASHIELD (Vector Laboratories) was used to mount coverslips. Slides were imaged using Leica fluorescence microscope. For staining of cultured cells, 4% PFA was used to fix cells for 15mins, followed by wash in PBS and blocking buffer. Cells were incubated with primary antibodies overnight at 4°C**.** Next day, species-appropriate goat/donkey secondary antibodies coupled to AlexaFluor dyes (488, 568 or 647, Invitrogen) and Hoechst dye for nuclear staining for 2 hrs at RT **or** overnight at 4°C**.** Slides were imaged using Leica or EVOS microscope.
5. **Proliferation assay:** GBM cells or TEC were plated at a density of 5000 cells per well in triplicates in 96-well plates. Proliferation was assessed 3 days after treatment with conditioned media or recombinant growth factors using CellTiter-Glo® Luminescent Cell Viability Assay. Luminescence signal was measured in a luminometer, and readings were taken on Day 0 of plating and at Day 3 or Day 5 after treatment to normalize for plating density and obtain the fold change in growth of cells.
6. **Western blotting:** Protein lysates were obtained from 300-500,000 cells per sample using RIPA buffer supplemented with protease and phosphatase inhibitors. Primary antibody for ASPN (Goat ab, Novus Biologicals, NB100-1514SS) and β-Actin (Rabbit mAb, Cell Signaling technology, 4970L) or GAPDH (mouse mAb) was used as loading control and incubated overnight at 4C. Secondary antibodies (anti-rabbit IgG HRP, Cell Signaling Technology and anti-goat HRP, Invitrogen) were added and incubated at room temperature for 2 hours. Protein bands were developed using the Clarity^TM^ western ECL substrate (Bio-Rad). Protein bands were visualized using MYECL gel imager (Thermo Scientific). Quantification of protein levels were done using Image J.
7. **Spheroid assay:** IDH1-mutant GBM cell line HK_252 was dissociated into single cells using Accumax. 4000 cells were seeded into each well of a low-attachment 96 well plate. The plate was then centrifuged at 600g for 5 minutes, and incubated at 37°C for 48 hours to allow the cells to aggregate and form individual spheroids. After spheroid formation, plate was imaged for initial area measurement (Day 0). Individual spheroids were then treated with either GBM media or TEC conditioned media. B27+ Heparin/EGF/FGF (HEF) mix was added to each TEC condition media to match the same concentration of B27+HEF as in the GBM media. The spheroids were incubated in conditioned media and cultured for 4 days, and imaged using LEICA microscope. The area of each sphere was calculated using ImageJ.
8. **Quantitation of intracellular D-2HG content**: An enzymatic D-2HG assay was used to measure the intracellular D-2HG content in TEC^26^. Briefly, cell pellets from LGG- and HGG-TEC lines were harvested in lysis buffer. The cell lysate was then divided into two parts: one for protein content determination by Pierce BCA protein assay kit (Thermo Scientific), and another for D-2HG measurement. For D-2HG assay, the lysate was first deproteinized by adding 3μl of Proteinase K (Qiagen) and incubated for 3 hours at 37°C, following which 25μl of lysate was added to 75μl of assay solution at RT for 30 minutes. Each sample was run as triplicates. Fluorometric detection was performed with Wallace Victor2 1420 Multi-label HTS Counter (PerkinElmer) at Em540nm/Ex610nm. D-2HG content was calculated based on a standard curve obtained from a range of concentrations of D-2HG replacing the cell lysate, and then expressed as D-2HG pmole/μg protein.
